# Genome-wide identification and expression analysis of the *FAR* gene family in hexaploid wheat (*Triticum aestivum* L.)

**DOI:** 10.1101/424739

**Authors:** Shandang Shi, Guaiqiang Chai, Hui Ren, Wenzhi Nan, Hongqi Wu, Yong Wang, Chunlian Li, Zhonghua Wang

## Abstract

Fatty acyl-CoA reductase (FAR) is involved in the biosynthesis of primary alcohols, which are waxy constituents that play an important role in plant stress. Previous studies have shown that primary alcohol is the most important component in the wheat seeding stage and accounts for more than 80% of the total composition. To date, eight *FAR* genes have been identified in wheat, but there has not been a systematic analysis. In this study, a comprehensive overview of the *TaFAR* gene family was performed, including analyses of the phylogenetic relationship, the multiple sequence alignment, the conserved motif distribution and the expression pattern. The result showed that a total of 41 wheat *FAR* genes were identified and designated *TaFAR1-A–TaFAR22-D;* all FAR genes were divided into six classes according to their phylogenetic relationship, and most of the *FAR* genes might be related to wheat cuticular wax synthesis. The analysis of the promoter binding site showed that *TaFAR* genes could be regulated by the MYB transcription factor and could be used as target genes for hormone regulation under adverse conditions, especially during a drought. This study provides a basis for further analyses of the *TaFAR* gene function and of upstream regulatory genes.

## Introduction

Wheat (*Triticum aestivum*) is one of the world’s most important food crops and feeds one-fifth of the population. Wheat yield is constrained by many factors, including biotic and abiotic stresses [1]. Drought is a major threat to wheat production [2]. The surface of wheat is covered with cuticular wax, which plays important roles in drought tolerance by limiting nonstomatal water loss [3]. Cuticular wax is a complex mixture of lipids and consists of very-long-chain fatty acids (VLCFAs) and their derivatives, including aldehydes, alkanes, alcohols, wax esters and ketones [4-7]. At the wheat seeding stage, primary alcohol is the most important component of cuticular wax and accounts for more than 80% of the total composition. Previous studies demonstrated that primary alcohols were synthesized by fatty acyl-CoA reductase (FAR) [8-13]. In *Arabidopsis*, the gene family contains eight members, and only the *AtFAR3/CER4* gene was involved in the primary alcohols biosynthesis of cuticular wax. However, eight *TaFAR* genes that are related to the biosynthesis of cuticular wax were identified in wheat, which suggested that there are a series of *TaFAR* genes involved in wax biosynthesis in wheat. With the gradual improvement of whole-genome sequencing and the annotation of wheat, it was possible to discover its *FAR* gene family [14].

A typical FAR protein contained an NAD(P)H binding Rossmann-fold (NADB) domain with Pfam ID: PF07993 and a fatty acyl-CoA reductase (‘FAR_C’) domain with PF03015 [15]. Thus, FARs were predicted to be extended short-chain dehydrogenase/reductase proteins at the N-terminus with an α/β folding pattern, a central β-sheet and a fatty acyl-CoA reductase domain at the C-terminus. All plant FARs contained two conserved motifs: the TGXXGXX(G/A) motif, which is involved in binding of NAD(P)H, and the YXXXK active site motif, which falls into the SDR117E family, which is a short-chain dehydrogenase/reductase superfamily [16-18]. In addition, because the first cloned *FAR* gene from *A. thaliana* encoded the MALE STERILITY2 (FAR2/MS2), the FAR_C domain was often annotated in databases as the “male sterile” domain. However, this annotation was outdated, because only two proteins in *Arabidopsis* and rice, At FAR2/MS2 and OsFAR2/DPW, affect male fertility [19, 20].

The *FAR* gene is also involved in the biosynthesis of suberin polyester and plant pollen development in addition to the synthesis of waxy components [21, 22]. Different *FAR* genes generally showed different functions according to the synthesized acyl chain lengths [7]. *AtFAR2/MS2* synthesizes primary alcohol during the stage of pollen exine development [23, 24]. *AtFAR3/CER4* was involved in the formation of C24:0-C30:0 primary alcohols in the cuticular wax of aerial organs [25]. *AtFAR1*, *AtFAR4* and *AtFAR5* generate 18:0, 20:0 and 22:0 fatty alcohols, respectively, which are present in root wax and suberin polyester [22]. In *Monocotyledonous*, partial genes of wheat, *Brachypodium distachyon* and rice had been identified. *OsFAR2/DPW*, an orthologous gene of *AtFAR2/MS2* in rice, affects the development and fertility of pollen. The expression of *OsFAR2/DPW* led to the formation of C16:0 primary alcohol by combining the substrates of C16:0 fatty acyl-CoA [26]. Three *FAR* genes had been identified in *Brachypodium distachyon*. The heterologous expression of *BdFAR1* results in the formation of C22:0 primary alcohol in yeast, while the expression of *BdFAR2* and *BdFAR3* led to the production of C26:0 primary alcohol [21]. In wheat, the function of eight *FAR* genes had been elucidated. The expression of *TaFAR1* and *TaFAR5* could produce C22:0 fatty alcohol in yeast and C26:0, C28:0, and C30:0 in tomato leaves. *TaFAR5* was also identical to *TaAA1b* as an anther-specific gene [1, 18]. *TaFAR2*, *TaFAR3* and *TaFAR4* were involved in the formation of C18:0, C28:0 and C24:0 fatty alcohols in yeast, respectively. *TaFAR4* was also identical to *TaAA1c* [17]. *TaFAR6* and *TaFAR8* catalyze the synthesis of C24:0, while *TaFAR7* synthesizes C26:0 in yeast [14]. *TaFAR6* was also identical to *TaMSF_2* as anther male sterility gene [27]. In addition, *TaMSF_1*, an anther male sterility gene, and *TaAA1a*, a fatty alcohols synthesis gene, were identified [1, 27]. In general, 8 *FAR* genes in wheat were characterized, but the information of the whole family is still unknown.

In this study, 41 *FAR* gene family members were identified from the bread wheat genome, and the detailed information on sequence homology, the phylogenetic relationship, the promoter analysis, and the expression patterns in various tissues were analyzed. This will be useful for further systematic functional characterization of wheat FAR genes.

## Materials and methods

### Plant materials and experimental design

Hexaploid wheat Chinese Spring (CS) was grown in a greenhouse of Northwest A & F University. The tissue samples were harvested at different stages, including leaves at 28-d-old, root at 28-d-old, spike at 39-d-old and stem at 65-d-old. Each sample was collected in at least three replicates, were quickly frozen in liquid nitrogen and were immediately stored at –80°C for further use.

### Identification of the *TaFAR* gene family

Two methods were used to identify wheat *FAR* genes. First, the protein database file of the whole wheat genome was downloaded from *EnsemblPlants* (http://plants.ensembl.org/index.html). A local BLASTP search was performed using *Arabidopsis* FAR proteins as queries against the wheat protein database with an e-value of e^−10^. Second, the ID of the conserved domain ‘PF07993’ was used to search genes in the *EnsemblPlants* database. Then, redundant candidates obtained by the two search methods were removed. An InterProScan Sequence Search (http://www.ebi.ac.uk/interpro/) was used to determine the presence of the FAR domain. Information on the coding sequence and the protein sequence was also obtained from the *EnsemblPlants* database. The Compute/Mw tool (http://web.expasy.org/compute_pi/) was used to predict the isoelectric point (pI) and the molecular weight (MW) of the wheat FARs [28].

### Sequence and conserved domain analysis of *TaFAR*s

The ClustalX program was used for multiple sequence alignments with default parameters [29]. Then, protein sequences were used to examine the conserved domain using a CD search (https://www.ncbi.nlm.nih.gov/Structure/cdd/wrpsb.cgi) and motifs using a MEME analysis online (http://meme-suite.org/tools/meme) with default settings [30, 31].

### Phylogenetic and promoter binding site analysis of TaFARs

A phylogenetic analysis was performed using MEGA 7 software through the method of neighbor-joining, and a bootstrap test was performed with 1000 replicates [32]. Two phylogenetic trees were produced: one contained only wheat FARs, and one used FAR proteins from wheat, *Arabidopsis*, rice, and *Brachypodium distachyon*. The promoter binding site was predicted by using 1500 bp upstream flanks of *TaFAR* genes in a *PlantCARE* database (http://bioinformatics.psb.ugent.be/webtools/plantcare/html/) [33].

### RNA-sequencing data analysis

The RNA-sequencing (RNA-seq) data of CS across the whole life cycle of wheat were downloaded from the WheatExp database (https://wheat.pw.usda.gov/WheatExp/). The transcript abundance of given genes was represented by the FPKM (fragments per kilobase of exon model per million mapped reads) values from diverse developmental processes. Heat maps of the *TaFAR* gene expression were generated using Cluster (http://soft.bio1000.com/show-119.html) and TreeView (http://soft.bio1000.com/show-17.html) software based on the FPKM values [34].

### Quantitative real-time PCR

The Plant RNA Purification Reagent (Invitrogen, USA) was used to extract the total RNA from each sample. The cDNA was synthesized using a PrimeScript reagent kit after treatment with RNase-free DNase I (Takara) according to the manufacturer’s instructions. The final cDNA samples were diluted 10-fold and were stored at –20°C for further use. For normalizing the gene expression in different RNA samples, the wheat *ACTIN* gene was used as an internal control [14]. The primers were designed using Primer Premier 5 (http://soft.bio1000.com/show-102.html) software (Table S1). The expression level of *TaFAR* genes was measured by quantitative real-time PCR (qRT-PCR) using a Bio-Rad Real-Time System (CFX96). Two independent biological repeats and three technical repetitions were produced and the quantification analysis was performed as described by Ma and Zhao [35].

### Data availability

The authors affirm that all data necessary for confirming the conclusions of the article are present within the article, figures, and tables. Supplemental material available at Figshare: <u>10.6084/m9.figshare.7110650</u>.

## Results

### Identification and sequence analysis of *TaFAR* genes

By searching in the database and submitting sequences to InterProScan, 41 *TaFAR* genes were finally identified. According to their phylogenetic relationship, those *TaFARs* family members were grouped into 22 clusters named *TaFAR1-A* to *TaFAR22-D* (Table 1, Figure 1A). Among them, 13 clusters were assigned to various A, B or D subgenomes. Those clusters were considered to be homoeologous copies of one *TaFAR* gene. The *TaFAR* gene information is listed in Table 1 and included the gene name, sequence accession number, protein length, MW and pI.

**Figure 1.**
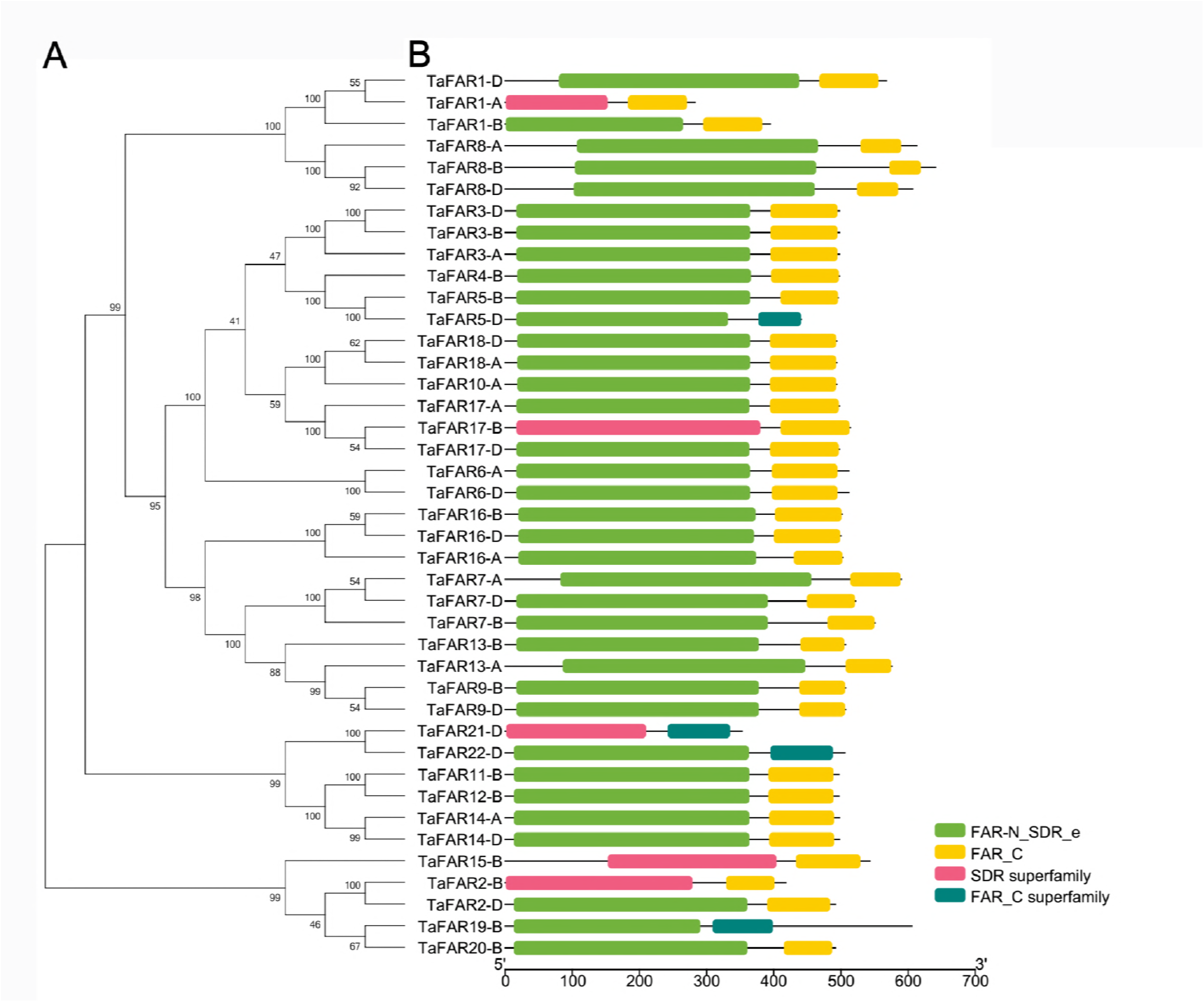
Phylogenetic analysis and the conserved domain of *TaFAR* genes and proteins. (A) Phylogenetic relationships of TaFAR proteins. The phylogenetic tree was produced using MEGA 7 software with the neighbor-joining method and bootstrap values were from 1000 replicates. (B) Two domains were found: FAR-N_SDR_e or SDR superfamily and FAR_C or FAR_C superfamily.

### Sequence analysis of TaFAR proteins

A multiple sequence alignment was performed using the amino acid sequences of TaFARs, which suggested that the TaFAR proteins contained two conserved motifs, the NAD(P)H binding site motif and the active site motif (Figure S1). The result of the conserved domain analysis showed that all TaFAR proteins contained conserved domains: FAR-N_SDR_e and FAR_C (Figure 1B). Among them, the SDR superfamily and FAR-N_SDR_e and the FAR_C superfamily and FAR_C had very similar functions according to the description on the InterProScan Sequence Search website. MEME analysis divided the protein sequence into 8 motifs (motifs 1–8), which are represented by different colors (Figure S2B). The FAR proteins are arranged according to the phylogenetic tree (Figure S2B). In addition, the extent of the conservation of every motif is represented by the height of each character by the online MEME (Figure S2A). The result of the sequence analysis showed that all TaFAR proteins are highly conserved.

### Phylogenetic analysis of genes in wheat

To further investigate the phylogenetic relationships between wheat, *Arabidopsis*, rice, and *Brachypodium distachyon*, a phylogenetic tree was constructed by aligning the protein sequences of 41 TaFARs, 8 AtFARs, 6 Bd FARs and 8 OsFARs. There were 65 FARs divided into seven groups named Classes 1,2a, 2b, 3–6. It is noteworthy that Class 2b contained only six members from *Arabidopsis*. The asterisk-tagged genes had been identified in wheat previously [5, 12]. The result showed that Classes 1, 3, 4, and 5 contain asterisk-tagged genes, which suggested that 32 *TaFAR* genes could be involved in wax synthesis. This result provides a basis for us to infer the function according to the phylogenetic relationship (Figure 2).

**Figure 2.**
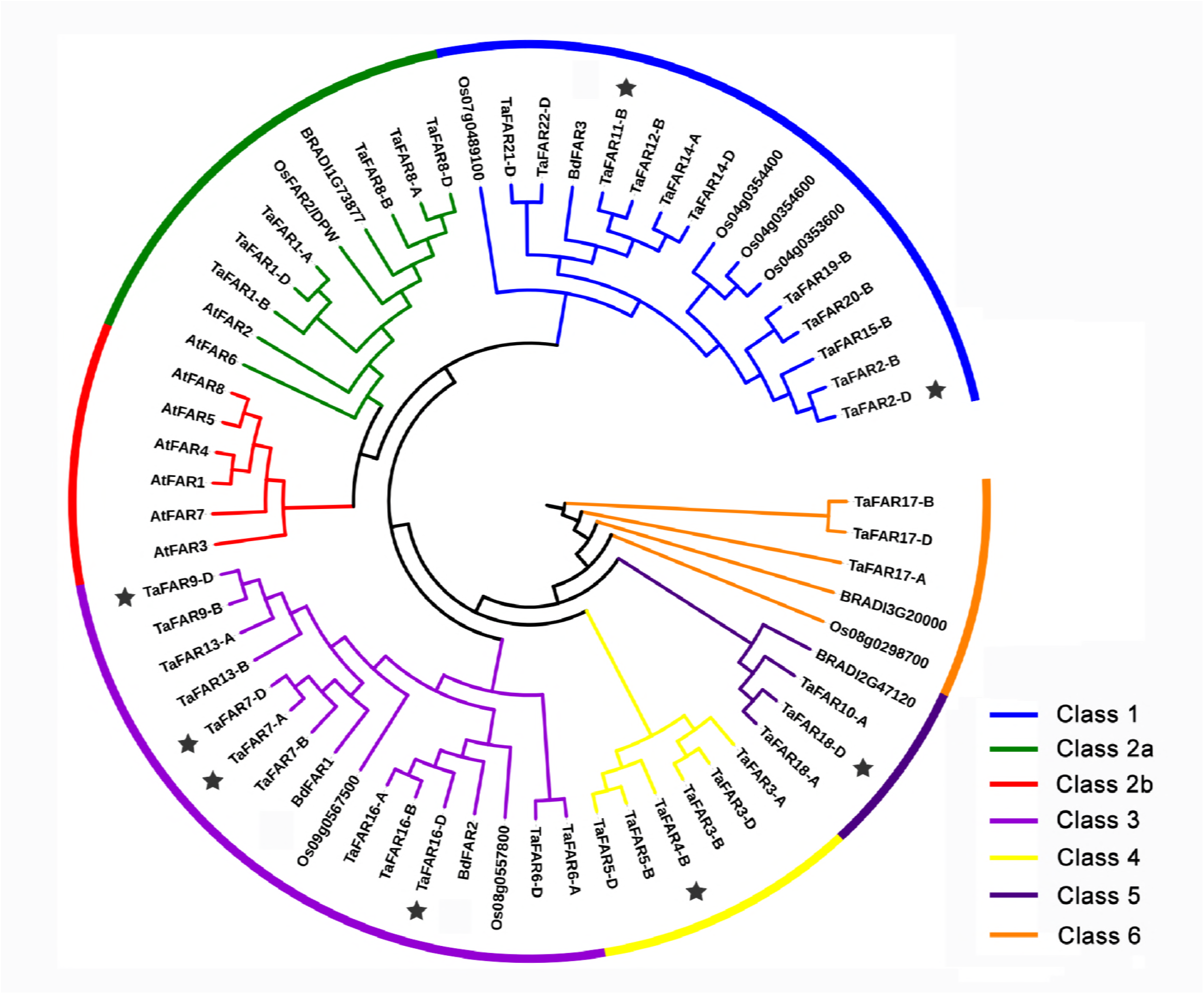
Phylogenetic analysis between wheat, *Arabidopsis*, rice, and *Brachypodium distachyon* FAR proteins. Phylogenetic relationships of wheat, *Arabidopsis*, rice, and *Brachypodium distachyon* FAR proteins. The phylogenetic tree was produced using MEGA 7 software with the neighbor-joining method and bootstrap values were obtained from 1000 replicates. These were divided into seven classes (Classes 1, 2a, 2b, 3-6), and Class 2b had no representative of wheat and contained only six members from *Arabidopsis*. The asterisk-tagged genes had been identified previously.

### Expression analysis of *TaFAR* genes in wheat

The transcript abundances of the *TaFAR* genes in various tissues are shown in Figure 3A. This result showed that almost all genes except *TaFAR8-A/B/D* have high expression levels during a certain period of the leaf. Most of the genes are also highly expressed in the stem and spike, and the expression of some genes, including *TaFAR8-A/B/D* in spike_z39, *TaFAR4-B/TaFAR1-B* in spike_z65, *TaFAR20-B* in stem_z65, are particularly high. Genes with a high expression in leaves, stem and spike could play an important role in plant cuticular wax synthesis. In addition, a small number of genes, including *TaFAR3-A/B/D* and *TaFAR5-B/D*, have high expressions in roots, and these genes may be related to the synthesis of suberin polyester (Figure 3A). Furthermore, a qRT-PCR analysis was used to test the consistency with the RNA-seq dataset. We randomly selected six genes to detect the expression levels of four tissues that correspond to the RNA-seq data. The results showed good consistency between the RNA-seq and the qRT-PCR data (Figure 3B).

**Figure 3.**
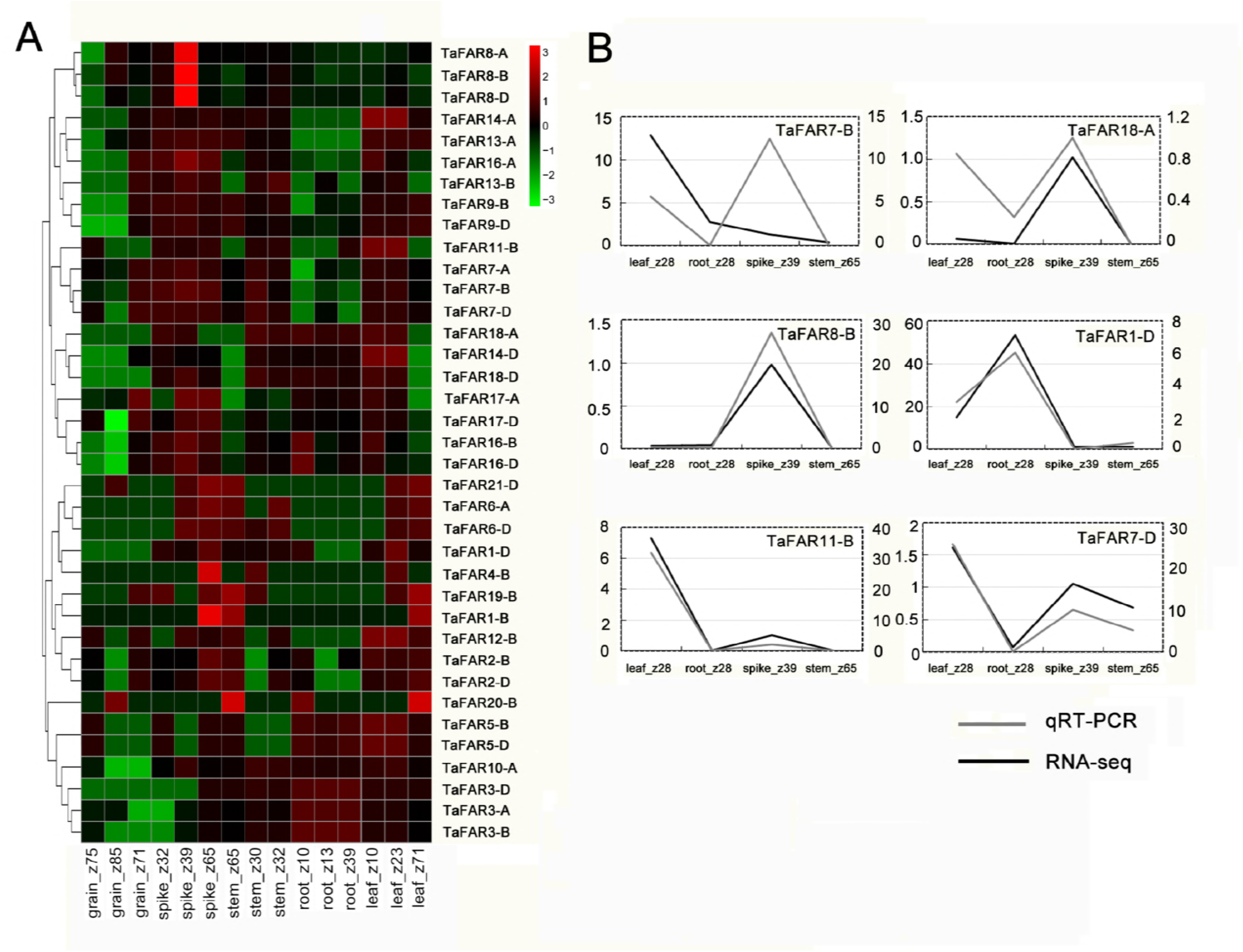
Expression profiles and QRT-PCR analysis of *TaFAR* genes. (A) Expression profiles analysis in different organs and tissues, including root, stem, leaf, spike and grain, with three difference stages, respectively. grain_z35 refers 35-d-old grain. The color scale represents different transcript abundances from low (blue) to high (red). (B) The consistency analysis of RNA-sequencing and qRT-PCR with regard to six *TaFAR* genes.

### Promoter binding site analysis of *TaFARs*

In this study, we selected 12 sites that are associated with the stress response from a number of promoter *cis*-regulatory elements and demonstrated their distribution in the *TaFARs* gene family (Figure 4A). The information on promoters is shown in Figure 4B. Five sites had wide distribution in the *TaFARs* gene family. The MYB binding site involved in drought-inducibility (MBS) was distributed to all members, suggesting that *FAR* genes are regulated by the MYB transcription factor under drought stress. For three hormone-related sites, including the cis-acting element involved in abscisic acid responsiveness (ABRE), the cis-acting regulatory element involved in MeJA-responsiveness (CGTCA-motif), and the cis-acting element involved in salicylic acid responsiveness (TCA-ELAMENT), almost all genes except *TaFAR14-A* contain at least one site, which suggests that *FAR* genes can respond to stress by binding hormone-including abscisic acid (ABA), salicylic acid (SA) or methyl jasmonate acid (MeJA). The cis-acting element involved in the defense and stress responsiveness (TC-RICH REPEATS) site is a cis-acting element that is involved in defense and stress responsiveness; its existence also proves that the *TaFARs* gene could respond to stress. Under heat treatment conditions, we analyzed the expression levels of eight genes that contain a cis-acting element involved in the heat stress responsiveness (HSE) site. The result showed that these genes had a consistent trend, in that the expression level decreased after one hour of treatment and increased after six hours of treatment (Figure 4C).

**Figure 4.**
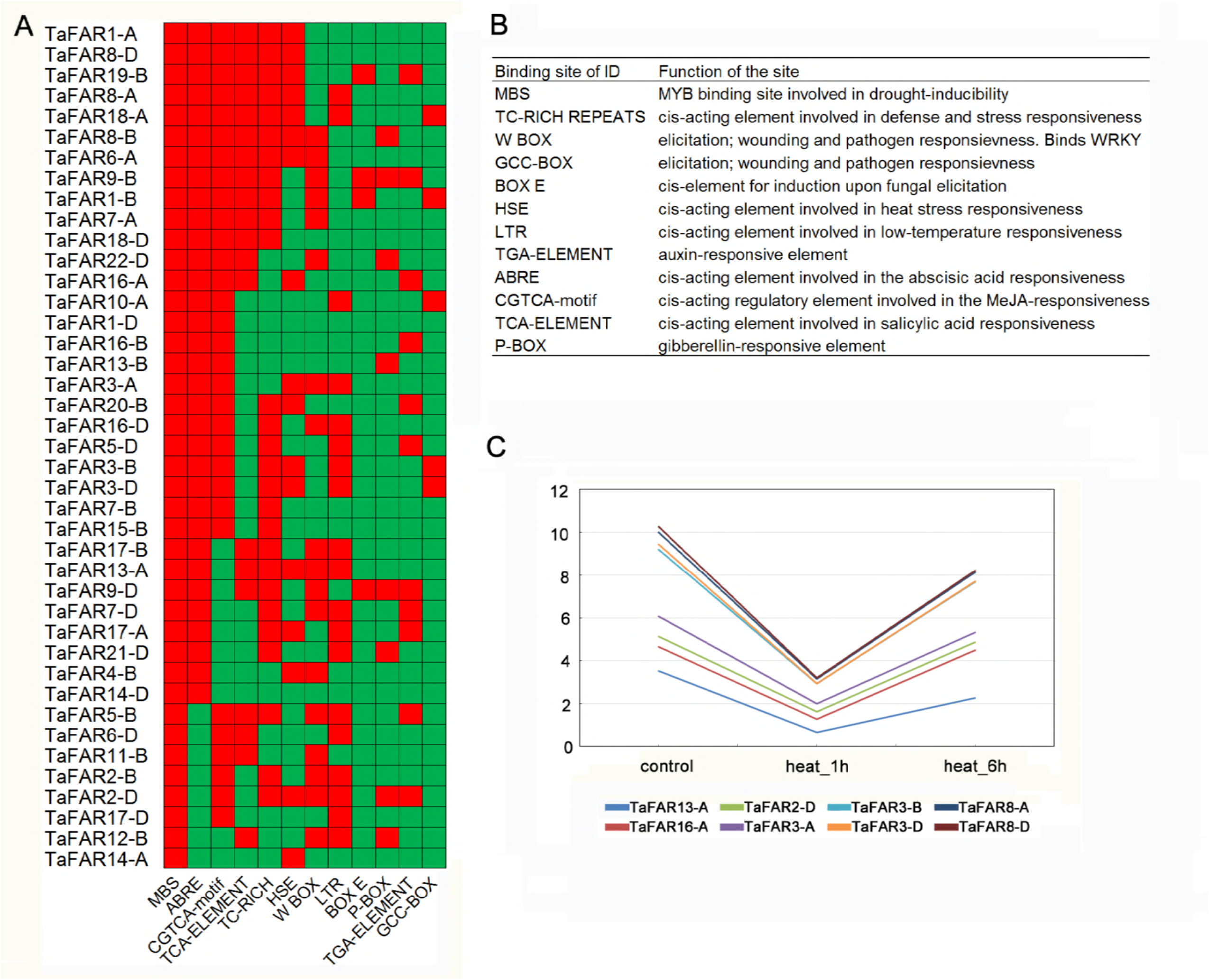
Promoter binding site analysis of *TaFARs.* (A) Distribution of putative *cis*-regulatory elements in 1500 bp upstream regions of 41 *TaFAR* genes. Red means there is this binding site; green means there is no such site. (B) Information on the promoter binding site. (C) Analysis of the expression levels of eight genes containing HSE under heat treatment conditions, including 0 h (control), 1 h and 6 h.

## Discussion

Wax is one of the important substances for plant drought resistance [3]. In the last few years, studies indicated that wax plays important roles in preventing the nonporous loss of water in wheat leaves [36]. The *FAR* gene family plays a critical role in water retention and stress response by synthesizing primary alcohols, which is a component of cuticular wax and accounts for more than 80% of the total composition at the wheat seeding stage [1]. Eight *FAR* genes were previously reported in wheat; they play an important role in the wax synthesis of wheat leaves [1, 18, 14, 17]. In *Arabidopsis*, only the *AtFAR3/CER4* gene was involved in the primary alcohols synthesis of cuticular wax, yet there are eight members in the *FAR* gene family [25]. These results suggest that there are a large number of unknown *FAR* genes in wheat. In this study, 41 *FAR* genes were identified in wheat and all of the *FAR* genes, including *Arabidopsis*, rice, wheat and *Brachypodium distachyon*, were divided into seven classes according to the phylogenetic relationship. The eight genes that had been identified as being related to wax synthesis were distributed in Classes 1, 3, 4, and 5. In addition, all of the genes in these classes have high expression levels in leaves, stem or spike (Figure 3A); thus, we could infer that there might be 32 *TaFAR* genes involved in the primary alcohols synthesis of wheat wax. In Class 2a, *AtFAR2* and *OsFAR2/DPW*, which are two male fertility genes, affected the development and fertility of pollen [19, 20], and we could speculate the possible functions of *TaFAR8-A/B/D* and *TaFAR1-A/B/D* in male fertility. In addition, *AtFAR1*, *AtFAR4* and *AtFAR5* had high expressions in roots and were involved in the synthesis of suberin polyester [22], which indicates that these genes, including *TaFAR3-A/B/D* and *TaFAR5-B/D*, may be related to the synthesis of suberin polyester. These functional predictions need to be proven by further experiments.

Wheat production was threatened by abiotic and biotic stresses [1]. Transcription factors and hormones are very important ways in which wheat plants respond to stress conditions. Current research showed that MYB transcription factors could regulate wax synthesis under drought conditions [37]. Plants could also resist drought by synthesizing the hormones of ABA, MeJA and Jasmonic acid JA [38, 39]. In this study, we analyzed the promoter binding site of *TaFARs* genes. Interestingly, all genes have an MBS site, which is the MYB binding site that is involved in drought-inducibility. The result showed that the *TaFAR* genes were indeed involved in drought resistance under the regulation of the MYB transcription factor, but the specific genes involved in regulation need further exploration. In our study, almost all of the *TaFAR* genes were contained in the hormone response sites of ABA, MeJA or JA. The result revealed that the *TaFAR* genes played an important role as target genes of ABA, MeJA or SA. In general, under drought conditions, the *TaFAR* genes were not only involved in the synthesis of cuticular wax by the regulation of MYB transcription factors but could also be used as target genes for hormones, including ABA, MeJA or SA, to resist drought. The wide distribution of the TC-RICH REPEATS site in the promoter region of the TaFAR gene family also suggested that *TaFAR* genes could be involved in response to other kinds of stresses. In addition, the expression levels of eight *FAR* genes containing HSE sites showed the same trend under heat stress. This trend of first decline and then rise indicated that the *FARs* were downregulated genes under heat stress, and this process might be a negative feedback regulation. This study provided important information for our next study to find upstream regulatory genes.

## Acknowledgments

This study was supported by National Natural Science Foundation of China (31471568).

---

**Figure S1 Multiple sequence alignment of TaFAR proteins.** A multiple sequence alignment was performed using the ClustalX program with default parameters. Two conserved motifs were marked by a horizontal line: the NAD(P)H binding site motif (TGXXGXXG) and the active site motif (YXXXK), where X represents any amino acid.

**Figure S2 Conserved motifs of TaFAR proteins.** (A) Compositions of the conserved motifs of TaFAR proteins. The extent of conservation of amino acid identity was represented by the height of each character. (B) The motif distribution of wheat, *Arabidopsis*, rice, and *Brachypodium distachyon* FAR proteins was investigated using the MEME web server. The FAR proteins were arranged according to the phylogenetic tree.

**Table 1. The *FAR* gene family in wheat (*Triticum aestivum*. L)**

**Table S1. Sequences of primers used in cloning and PCR reactions.**

